# Encompassing view of spatial and single-cell RNA-seq renews the role of the microvasculature in human atherosclerosis

**DOI:** 10.1101/2023.12.15.571796

**Authors:** Tore Bleckwehl, Sidrah Maryam, Anne Babler, Michael Nyberg, Markus Bosteen, Maurice Halder, Charles Pyke, Henning Hvid, Louise Marie Voetmann, Judith C. Sluimer, Vivek Das, Simon Baumgart, Rafael Kramann, Sikander Hayat

## Abstract

Atherosclerosis is a pervasive contributor to cardiovascular diseases including ischemic heart disease and stroke. Despite the advance and success of effective lipid lowering-therapies and hypertensive agents, the residual risk of an atherosclerotic event remains high and improving disease understanding and development of novel therapeutic strategies has proven to be challenging. This is largely due to the complexity of atherosclerosis with a spatial interplay of multiple cell types within the vascular wall. Here, we generated an integrative high-resolution map of human atherosclerotic plaques by combining single-cell RNA-seq from multiple studies and novel spatial transcriptomics data from 12 human specimens to gain insights into disease mechanisms. Comparative analyses revealed cell-type and atherosclerosis-specific expression changes and associated alterations in cell-cell communication. We highlight the possible recruitment of lymphocytes via different endothelial cells of the vasa vasorum, the migration of vascular smooth muscle cells towards the lumen to become fibromyocytes, and cell-cell communication in the plaque, indicating an intricate cellular interplay within the adventitia and the subendothelial space in human atherosclerosis.

## Main

Atherosclerosis, a complex disease with many clinical manifestations, is a leading cause of morbidity and mortality worldwide^1^. Atherosclerosis refers to the accumulation of fatty and/or fibrous material in the intima of the arterial wall. With time, the atherosclerotic plaque can become more fibrous and accumulate calcium^2^. The accumulation of these substances results in a complex pathophysiology with various cellular interactions and modulations involved that can result in vascular occlusion and/or plaque rupture or erosion that clinically manifests as myocardial infarction and stroke^3^. Importantly, there is accumulating evidence supporting a high degree of diversity of mechanisms underlying atherosclerotic disease and related thrombotic events^4,5^. Recent single-cell sequencing studies have revealed cellular resolution insights into atherosclerotic tissue in mice and humans, shedding light on cellular heterogeneity, cellular plasticity, and the contribution of different cell types to atherogenesis^6^.

However, despite intensive preclinical research on atherosclerosis, a firm understanding of human pathogenesis is still lacking, and the current scRNA-seq studies are limited in patient diversity due to low sample numbers. In addition, independent studies might have technical batch effects that limit the comparability of the results. To overcome the challenges associated with a limited number of samples per study and technical batch-effects, we have created a comprehensive technical batch-effect corrected unified human atherosclerosis atlas. Furthermore, to account for spatial patterns and their association with progressive disease states, we generated a novel spatially resolved transcriptomics dataset with 12 human samples. Our integrated atlas with single-cell and spatial transcriptomics presents insights into structural and collective functions of cell types in their spatial niche of the human artery.

Integrative analyses of our novel atlas indicate the importance of pericytes and phenotypically distinct endothelial cell clusters in human atherosclerosis. Analysis of the spatially resolved human artery data led to the identification of lymphocyte accumulation in the adventitia and possible cell-cell-interaction with endothelial cells and pericytes of the microvasculature.

A pivotal characteristic in atherosclerosis lies in the intricate orchestration of lipoprotein uptake and efflux within the atherosclerotic plaque, primarily instigated by scavenger receptors predominantly found on macrophages^7^. With the spatial transcriptomics data we highlight major interactions of this process and highlight a possible lipoprotein affiniation of pericytes in the adventitia.

Furthermore, using computational pseudo-time trajectories, we found a potential transition of vascular smooth muscle cells (VSMC) into fibromyocytes from the outer medial layer towards the lumen and progression of phenotypical foamy cells into SPP1 positive macrophages, accompanied by reorganization of the extracellular matrix in the subendothelial layer. Finally, using a database of existing drugs, we showcase the potential of our atlas in aiding the identification of novel drug targets that are specific to relevant cell states and spatial niches. Overall, our findings point toward a collateral or initiating adventitis in atherogenesis and highlights avenues for further research.

## Results

### Functional analysis of the integrated human single-cell atherosclerosis atlas

To attain a better understanding of the cellular heterogeneity, and interactions between diverse cell types within human atherosclerotic plaques, we integrated single-cell RNA-sequencing (scRNA-seq) datasets encompassing coronary and carotid human arteries. Our data integration approach encompassed 4 studies involving 10 atherosclerotic donors^8–10^ and 3 control donors^11^, culminating in 130,000 high-quality single cells after a rigorous quality filtering process including doublet removal, and technical batch-effect correction. The integrated and manually annotated atlas revealed new insights into atherosclerosis and might serve as a valuable reference for forthcoming atherosclerosis studies and novel drug target identification. Furthermore, our atlas contains novel spatial transcriptomics data from different atherosclerosis stages of human coronary arteries (**Fig. 1a**).

**Figure 1:**
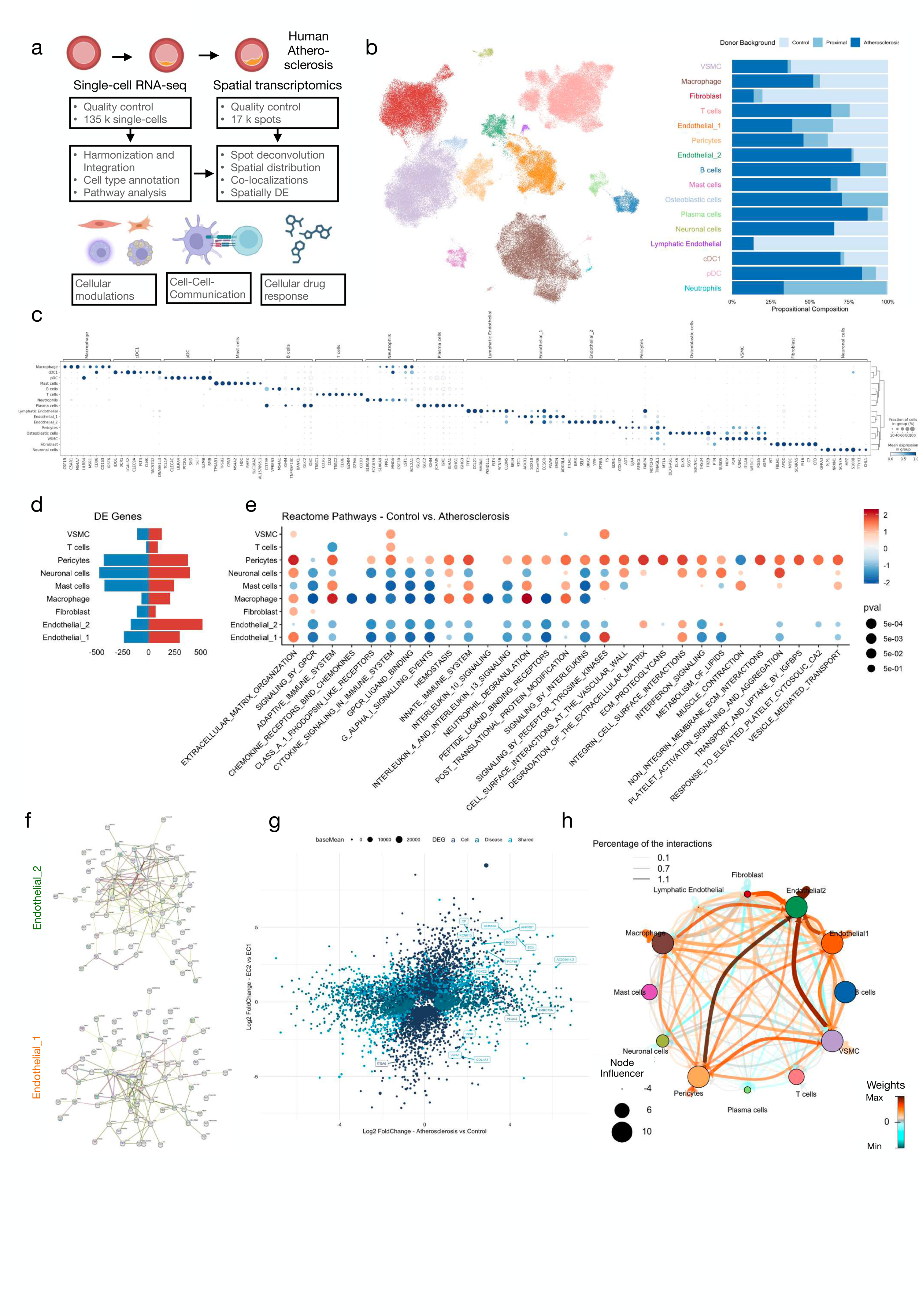
Functional analysis of the integrated human single-cell atherosclerosis atlas. a - Overview of data, quality control procedures, and analysis: Description of the single-cell RNA-seq and spatial transcriptomics data incorporated into the human atherosclerosis atlas, including the quality control (QC) process and analytical methodologies. b - UMAP embedding: Uniform Manifold Approximation and Projection (UMAP) visualization of the scVI-integrated single-cell dataset. Cell clusters were identified through Leiden clustering (resolution 0.2) and further annotated based on marker gene expression. Clusters are arranged by total cell count, with disease background proportions indicated. Proximal cells were isolated from regions adjacent to carotid artery plaques (Alsaigh et al.). c - Identification of top marker genes: Displaying key marker genes characterizing distinct cell types within various clusters of the human atherosclerosis single-cell RNA-seq atlas. d - Differential gene expression within main clusters: Number of genes identified within each cluster by differential expression analysis of the scVI model. e - Reactome pathway enrichment analysis: Comparative assessment of pathway enrichment between control and atherosclerosis conditions within major cell types. Color gradient denotes normalized enrichment scores (NES), while size indicates associated p-values derived from gene set enrichment analysis. f - Protein-Protein Interactions: Representation of protein-protein interactions among the top differentially expressed genes identified via *stringdb* for Endothelial_2 (top) and Endothelial_1 (bottom). g - Differential expression analysis of pseudo bulk: Comparison of atherosclerosis versus Control states (x-axis) across different endothelial clusters (y-axis) using *DESeq2* for pseudo bulk differential expression analysis. h - Ligand-Receptor interaction network: Network analysis generated by *CrossTalkR*, highlighting enriched cell-cell communication via ligand-receptor interactions within major cell types in human atherosclerosis.

For the construction of our human single-cell atlas for atherosclerosis, we curated and harmonized the data of diverse scRNA-seq datasets (**Extended Data Fig. 1a-b**) by conducting a comparative evaluation of different scRNA-seq integration technologies (**Extended Data Fig. 1c**). Here, Harmony^12^ and scVI^13^ emerged as particularly proficient integration tools for the single-cell human atherosclerosis atlas, with high donor mixtures and comparable cell type and disease compositions for both methods (**Fig. 1b, Extended Data Fig. 1d-e**). With the scVI-based embedding, we identified 16 main cell type clusters through marker gene expression profiling (**Fig. 1c, Extended Data Fig. 1f, Supplementary Table 1**). Among these clusters, we found a cluster of pericytes (e.g. COX4I2, HIGD1B) and osteoblastic cells (e.g. DLX5/6). Particularly intriguing, we found two distinct endothelial cell clusters, with specific and elevated expression of DKK2 and ACKR1 expression, respectively, but shared comparable expression of endothelial markers (e.g. VWF, PECAM1). Notably, cellular compositions of non-immune cell-types such as VSMCs and pericytes were balanced between control and atherosclerosis, while immune-related clusters such as B cells and T cells had more cells from atherosclerotic donors (Fig. 1b).

Having defined the main cell types, we analyzed the differential expression between control and atherosclerosis states in each cell type (**Fig. 1d, Supplementary Table 2**). Based on these differentially expressed genes, we performed pathway enrichment analysis for each cell type to identify any systematic patterns in gene dysregulation. Interleukin-10 signaling was enriched in the control cells of macrophages, consistent with its athero-protective influence^14^, while immune responses were enriched in atherosclerotic macrophages (**Fig. 1e**). Surprisingly, we identified a broad dysfunction in pericytes, which include a plethora of pathway changes in the extracellular matrix (ECM), IGFBP transport and immune cell-related responses (**Extended Data Fig. 1g**). The comprehensive transcriptome alterations of periycytes was somewhat surprising as their role in Atherosclerosis is understudied^15^. However, the relatively diminished gene expression profile observed in pericytes compared to other cellular counterparts, coupled with their transcriptional resemblance to vascular smooth muscle cells (VSMCs), potentially masked the distinctive impact of pericytes. Consequently, the revelation of perturbed genes and pathways specific to pericytes within our single-cell atlas underscores the significance of employing single-cell studies to discern cell-type-specific associations with diseases.

Notably, the transcriptional changes were impacted differently in the two different endothelial cell clusters which differed significantly in ECM remodeling. Differential expression analysis of the two endothelial cells showed that among other ECM components, COL15A1 (higher expressed in Endothelial_1) or COL8A1 (higher expressed in Endothelial_2) differ (**Extended Data Fig. 1h**). Furthermore, the two types of endothelial cells showed different potential gene interactions: Endothelial_1 gene interactions are associated with ITGA6 and EMCN (**Fig. 1f**), while the Endothelial_2 cluster showed expression of mesenchymal related transcription factors (FOXC2, TBX1). Taking into account the differential expression of the two types of endothelial cell clusters and their disease state, we revealed common (LRRC75A, PLCG2) and endothelial cluster specific atherosclerotic genes, such as COL4A1 and VWA1 for Endothelial_1 and the aforementioned mesenchymal transcription factors as well as SCX or KCNK15 for Endothelial_2 (**Fig. 1g**).

Finally, we analyzed the cell-cell communication of all cell types of the human atherosclerosis atlas with *CellPhoneDB^16^* and *CrossTalkerR^17^* to identify ligand-receptor pairs enriched in atherosclerosis compared to the controls. Among the 1,500 interactions analyzed from the LIANA database, we found an interactive network between pericytes, endothelial cells and VSMC as well as macrophages (**Fig. 1h**, **Supplementary Table 3**). Taken together, we integrated available single-cell RNA-seq data from human atherosclerosis and controls, identified two different endothelial cell clusters, transcriptional changes especially in the ECM, and an altered cell-cell communication network involving multiple cell-types in atherosclerosis.

### Spatial transcriptomics of human atherosclerosis

Given the new insights from the human atherosclerosis atlas, we next wanted to investigate their spatial distribution in human arteries and performed spatial transcriptomics of 12 human coronary arteries of different disease stages (**Fig. 2a**). In total we captured 16593 spots with 1563 genes per spot on average (**Extended Data Fig. 2a**). With an estimation of 5 cells per spot, we leveraged *cell2location^18^* to deconvolute the intricate cellular composition of the tissue slides with our scRNA-seq atlas as reference. Cell type abundances were well distributed across the arteries and the correlation of the reference signatures was low between different cell types, indicating a commendable capacity to deconvolute the spatial location of the cell types throughout the complete artery (**Extended Data Fig. 2b-c**).

**Figure 2:**
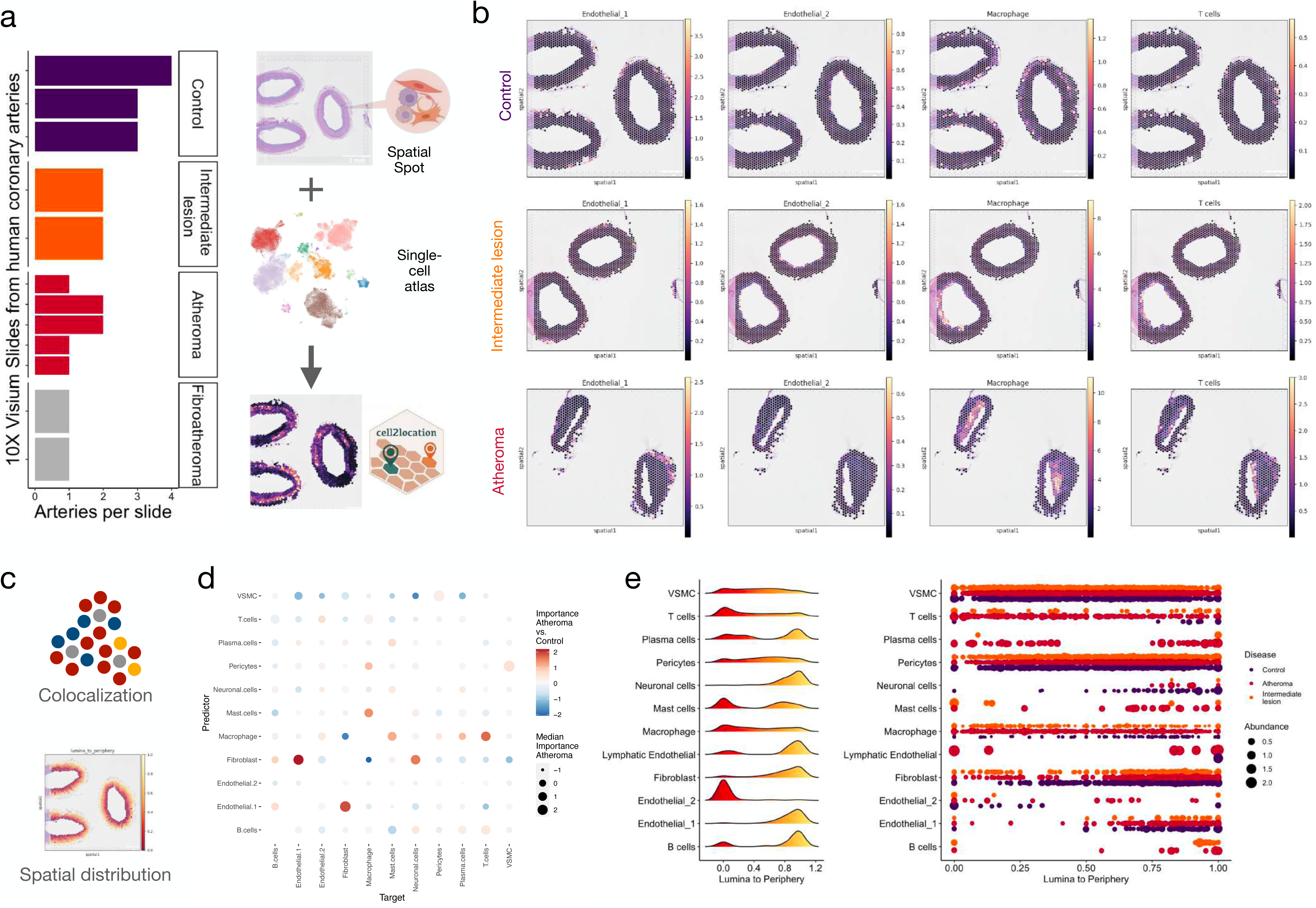
Spatial transcriptomics of human atherosclerosis. a - Study design and spatial transcriptomics overview: Depiction of the study design detailing the donor backgrounds across different 10X Visium slides obtained from human coronary arteries (left). On the right, a schematic representation illustrates Visium spot deconvolution for estimating cell type abundances using *cell2location* with the human scRNA-seq atherosclerosis atlas. b - Illustration of cell type abundances in Visium slides: Examples showcasing cell type abundances observed in Visium slides derived from specimens of control, intermediate lesions, or atheroma within the coronary artery. c - Visualization of co-localizations and Lumina-to-Periphery score calculation: Spots proximal to the lumen are assigned a value of 0, while spots on the outermost layer of the adventitia are assigned a value of 1. d - Co-occurrence of cell type abundance within and among spots: Within the same spot and between juxtapositional spots the co-occurrence was estimated with *misty* across all slides, contrasting atheroma slides versus control. e - Ridgeline Plot of Lumina-to-Periphery scores across control, intermediate lesion, and atheroma slides: Displaying the relative positioning of each cell type within the artery based on the lumina-to-periphery score for all slides. Additionally, the plot on the right separates the same data by disease background, with the size of the plot segments indicating normalized cell type abundances.

Interestingly, we observed that the majority of cell types exhibited a distinct and consistent positioning within the artery (**Fig. 2b, Extended Data Fig. 2d**). This was especially evident for most immune cells in the plaque, fibroblasts in the adventitia and VSMC in the media as expected. However, the two endothelial cell clusters were found to be located mutually exclusively in the luminal endothelium or adventitia, respectively. EC markers, which were expressed in both endothelial cell types (e.g. VWF) correlated with the abundance of luminal and peripheral endothelial cells, while cluster-specific markers obtained from differential expression analysis of the two EC types were located with the corresponding endothelial location (**Extended Data Fig. 3a**).

Next, we applied the Multiview Intercellular SpaTial modeling framework (MISTy)^19^ to analyze the neighborhood of each major cell type by predicting their colocalization to other major cell types within the same spot and the adjacent spots. By comparing the slides with control tissue and the slides with the atherosclerosis tissue, we found that VSMC and pericytes showed a general colocalization, but macrophages changed their association with fibroblasts to other immune cells in the atheroma slides (**Fig. 2c-d, Extended Data Fig. 3b**). On the other hand, Endothelial_1 could predict the localization of fibroblasts, and intriguingly to some extent even Lymphocytes. This was particularly interesting, as with the progression of atherosclerosis, T cells and macrophages were identified with high abundance in the plaque region of all atheroma slides, while Endothelial_1 were highly abundant in the adventitia.

To systematically evaluate the regionality of the deconvoluted cell types in our spatial coordinates, we defined a gradient from the most inner spot (tunica intima) of all arteries towards the adventitia (**Fig. 2c**). This approach showed that the spatial separation of the endothelial cell types was observed in all slides (**Fig. 2e**). Further, fibroblasts were mostly located in the periphery, in agreement with their association with the adventitia^20^. In contrast, immune cells followed a bimodal distribution and were located in the inner arterial wall as well as in the adventitia. The accumulation of the immune cells in both areas was more evident with the progression of atherosclerosis.

### Peripheral endothelial and pericytes are linked to lymphocyte adhesion

Given the identification of two different endothelial cells in the intima and the adventitia and a bimodal distribution of immune cells, we next asked if this would reveal a difference in immune cell adhesion and recruitment. When we compared the atherosclerosis associated Ligand-Receptor interaction scores of immune cells with the different endothelial cells, we found an increased cell-cell communication between peripheral ECs and pericytes with lymphocytes (**Fig. 3a**). This was observed irrespective of whether the immune cells act as ligand or receptor expressing cells. In particular, we found that Major Histocompatibility Complex (MHC II) genes were more abundantly expressed in peripheral endothelial cells, which could interact with immune cells by presenting antigens^21^. In addition, the counterparts of other ligands and receptors that were expressed in peripheral endothelial cells showed elevated expression levels in T-cells (**Fig. 3b**). Notably, the lymphocyte interaction partners on peripheral endothelial cells did not show elevated expression on lymphatic endothelial cells. Next, we performed spatial cell-cell-communication analysis^22^ for the ligand-receptor pairs defined between peripheral endothelial cells and T-cells in the spatial transcriptomics data, which showed potential interactions of the MHC II gene HLA-DBP1 and CD4 in peripheral endothelial sites as well as in the plaque region (**Fig. 3c**). The potential interaction in the plaque is consistent with high expression of MHC II in macrophage/cDC1 in general (**Extended Data Fig. 3c**). Furthermore, receptor expression in peripheral endothelial cells were more specific to these endothelial cells and hence cell-cell interactions were mainly observed in the adventitia at sites with high endothelial cell abundance. This is exemplarily shown for the interaction between CCL5 and ACKR1. In contrast, the immune cell related interactions of CCL5 with CCR1 were dominating the plaque core region.

**Figure 3:**
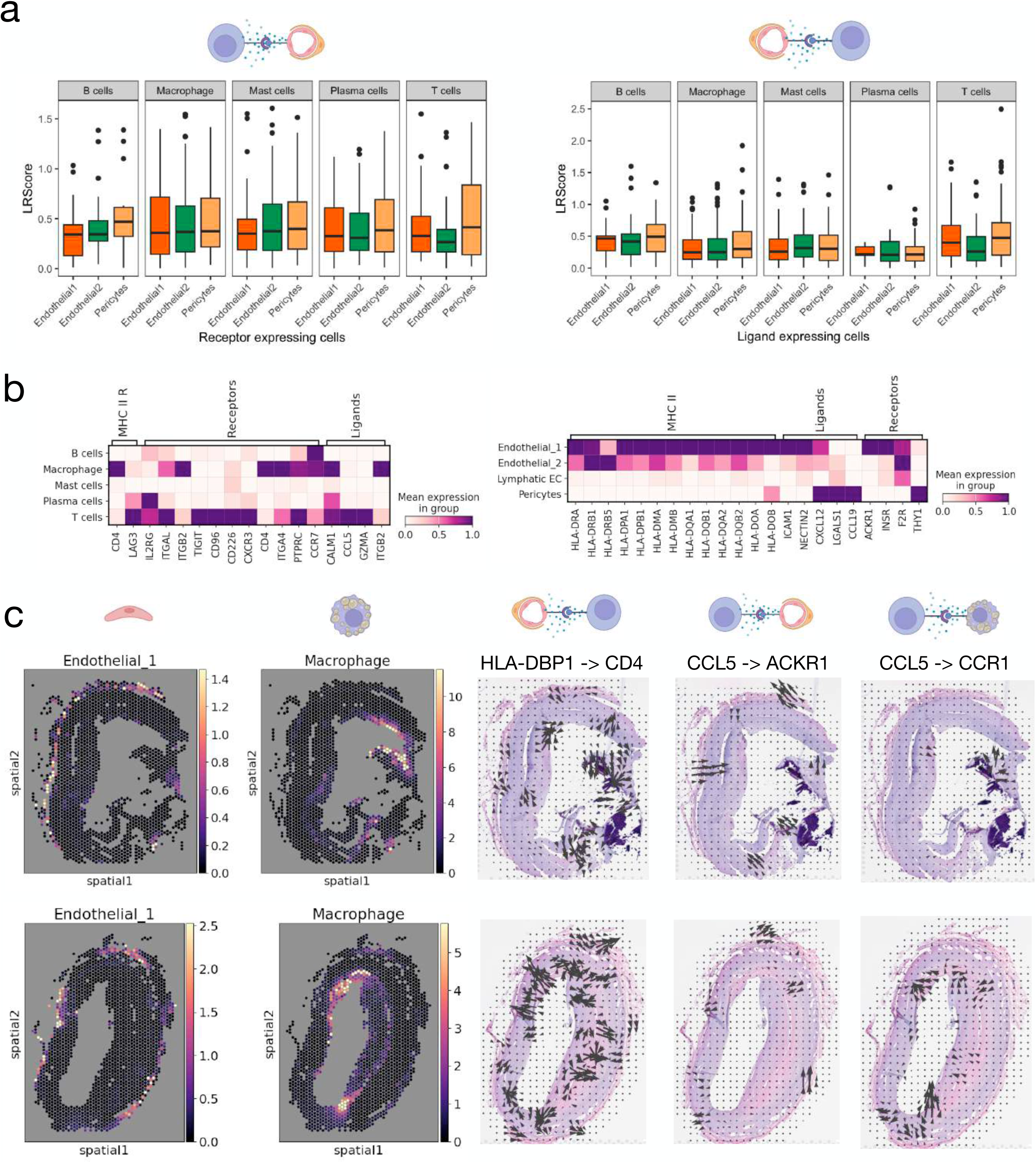
Peripheral endothelial and pericytes are linked to lymphocyte adhesion. a - Ligand-Receptor interaction between lumina and peripheral endothelial cells and immune cells: Utilizing data from the human atherosclerosis scRNA-seq atlas, the Ligand-Receptor Score (LRScore) demonstrates the weighted interaction score for atherosclerosis, highlighting interactions between luminal and peripheral endothelial cells along with immune cells. b - Expression of receptors and ligands in endothelial cells, pericytes, and immune cells: Presentation of diverse receptor and ligand expressions in various endothelial cell types, pericytes, and their counterparts across different immune cell subsets. c - Illustration of spatial ligand-receptor interactions: Displaying examples of spatial relationships in ligand-receptor interactions. The left side showcases the abundance of peripheral endothelial cells and macrophages within two atheroma slides. Examples of endothelial ligands paired with T-cell receptors (HLA-DBP1 -> CD4), T-cell ligand paired with endothelial receptors (CCL5 -> ACKR1), and T-cell interactions with macrophages (CCL5 -> CCR1) are depicted.

In summary, the ligand receptor analysis indicates potential adhesion and recruitment of lymphocytes via peripheral endothelial cells of the vasa vasorum that might be supported by pericytes which are known to have intricate connections with the endothelium of the vasa vasorum.

### Subclustering reveals cell type changes in the arterial microenvironment

To gain a deeper understanding of the human atherosclerosis atlas and our spatial transcriptomics data, we next performed subclustering of the single-cell RNA-seq data. Our sub-clustering revealed 32 cell states across all major cell-types. Specifically, we could differentiate fibromyocytes and myofibroblasts from VSMCs and two different pericyte clusters (APOE+ and MYH11+) as well as different macrophage and T-cell clusters (**Fig. 4a**). Within the T-cell cluster, we identified different T-cells and two types of natural killer cells, with one possibly being cytotoxic (GZMB, GNLY) and enriched for MHC II antigen presentation and interferon gamma and the other (XCR1, CD56/NCAM1, FCGR3A/CD16) expressing parts of the Toll-like receptor cascade (**Extended Data Fig. 4a-b**). For the macrophages, subclusters of monocyte-derived dendritic cells (CDC1 and FCER1A), Resident macrophages (LYVE1), inflammatory macrophages (IL1B and TNF), M4 macrophages (S1008/9), foamy macrophages (TREM2) and SPP1+ macrophages were annotated, whereby foamy macrophages showed enrichment in GPCR pathways and SPP1+ macrophage in ECM and lipid metabolism (**Extended Data Fig. 4c-d**).

**Figure 4:**
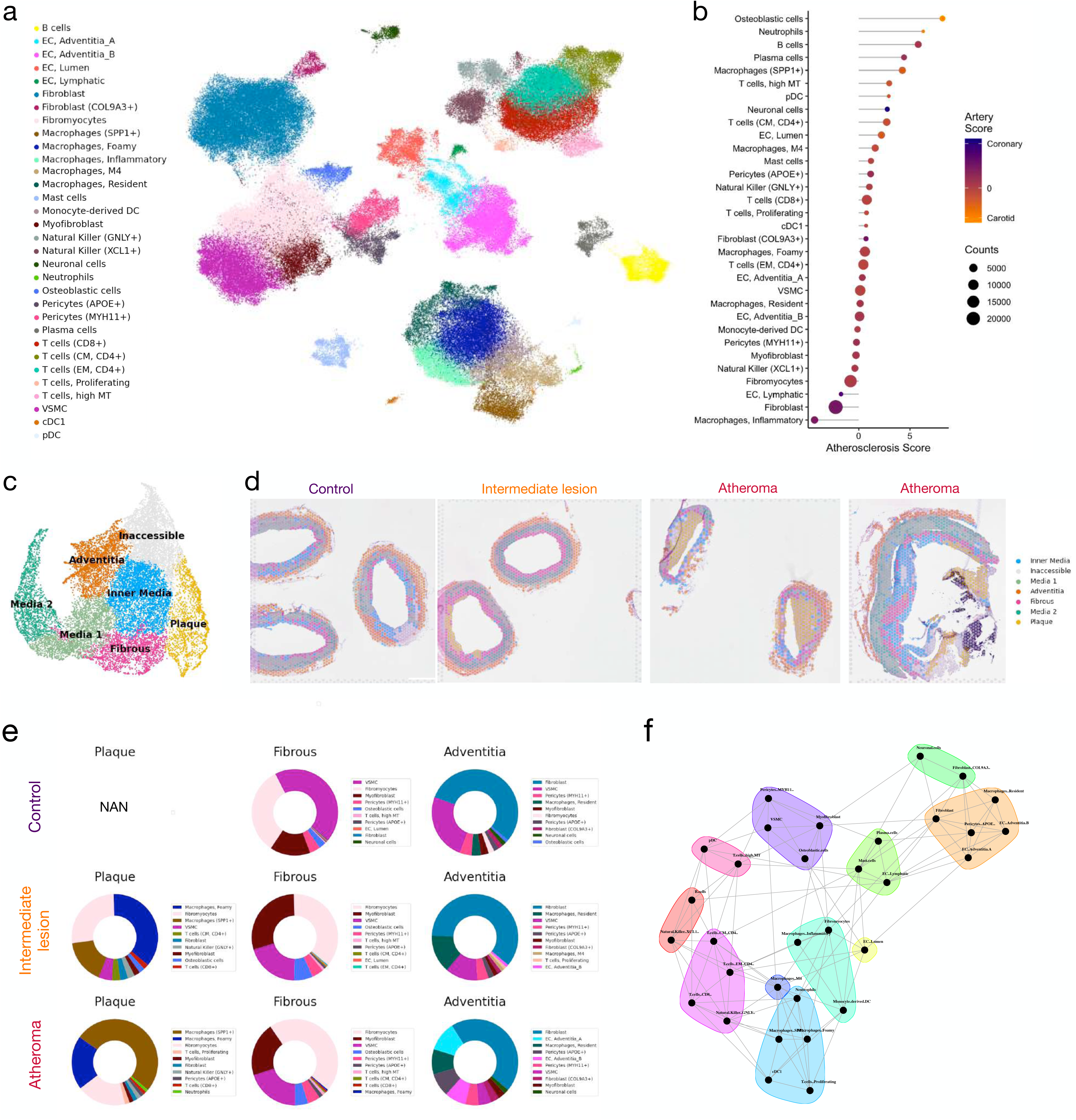
Subclustering reveals cell type changes in the arterial microenvironment. a - UMAP of scVI-Integrated scRNA-seq data with subclustering annotation: The major cell types were clusters separately and the resulting clusters were annotated by marker gene expression evaluation. b - Subclusters Ranking: Displaying the ranking of cell type subclusters according to their relative contributions to atherosclerosis. The x-axis denotes the log2 ratio of normalized cell counts between control and control state, while color indicates the ratio between sampling from coronary versus carotid arteries. Size correlates with the total cell count for each subcluster. c - UMAP of the Visium spots and their clustering by the cell type abundance: The spatial clusters were annotated according to their location within the slides or read coverage. d - Spatial clustering examples in representative Visium slides showcasing spatial clustering for specific cell types observed in samples from control, intermediate, and atherosclerotic coronary arteries. e - Median cell type abundance within the spots of each spatial microenvironment indicated for the sample from control, intermediate and atherosclerotic coronary arteries. f - Visualization of community interactions among deconvoluted cell types within atherosclerotic slides.

Based on the identified clusters, and accounting for study-size and disease status, we showed that osteoblastic and neuronal cell abundance increased in carotid and coronary artery atherosclerosis, respectively. (**Fig. 4b**). Beside these artery specific alterations, we found a general higher presence of leukocytes in both arteries as expected, but also observed an increased cell abundance in luminal EC and pericytes (APOE+).

Next, to investigate the spatial distribution of the defined cell types and their spatial association with atherosclerosis, we first clustered the spatial spots into microenvironments and termed these according to their location within the artery (**Fig. 4c-d**). The plaque and fibrous microenvironment clusters were mostly composed of spots from atherosclerotic specimens (**Extended Data Fig. 4e-f**) and within the plaque cluster, foamy macrophages were observed in the intermediate lesions with enrichment of SPP1+ macrophages with disease progression (**Fig. 4e, Extended Data Fig. 4g**). Starting with the intermediate lesions, the fibrous cluster was enriched for fibromyocytes and in the adventitia, the proportion of fibroblasts was skewed towards an increased proportion of peripheral EC, pericytes (APOE+) and resident macrophages, which might indicate microvascular angiogenesis^23^. Furthermore, the media clusters were dominated by a high abundance of VSMC, as expected (**Extended Data Fig. 4h**).

Finally, we analyzed the colocalization of the subclusters revealing complex interactions of the deconvoluted cell states (**Fig. 4f**), whereby APOE pericytes were found to be adjacent to cell types of the adventitia and MYH11 pericytes were close to media related cell types. Additionally, fibromyocytes were related to media and plaque cell clusters and lymphocytes were distinct from the plaque, indicating their bimodal role. Next, our aim was to study cell-cell communication leveraging spatial transcriptomic information to validate ligand-receptor interactions by proximity. Because the colocalization favors the interaction of highly expressed genes from the same cell type, the most relevant interactions of the atheroma slides were identified in the plaque (**Extended Data 5a, Supplementary Table 4**). However, we linked each ligand receptor pair to cell type expression and specificity of the scRNA-seq atlas, which revealed the spatial proximity of cell-cell interactions across different cell types that will be highlighted in the following sections.

### Lipid uptake, efflux and interplay of the macrophages in the human plaque

Before investigating the spatial relationships within the plaque, we first analyzed the spatial distribution of the different macrophages in the intermediate disease and atheroma slides. As expected, Foamy macrophages and SPP1 macrophages were highly accumulated towards the lumen (**Fig. 5a-b, Extended Data 5b**). M4 macrophages were located in the adventitia and plaque, consistent with reports from other human coronary arteries^24^, while resident macrophages dominated the adventitia.

**Figure 5:**
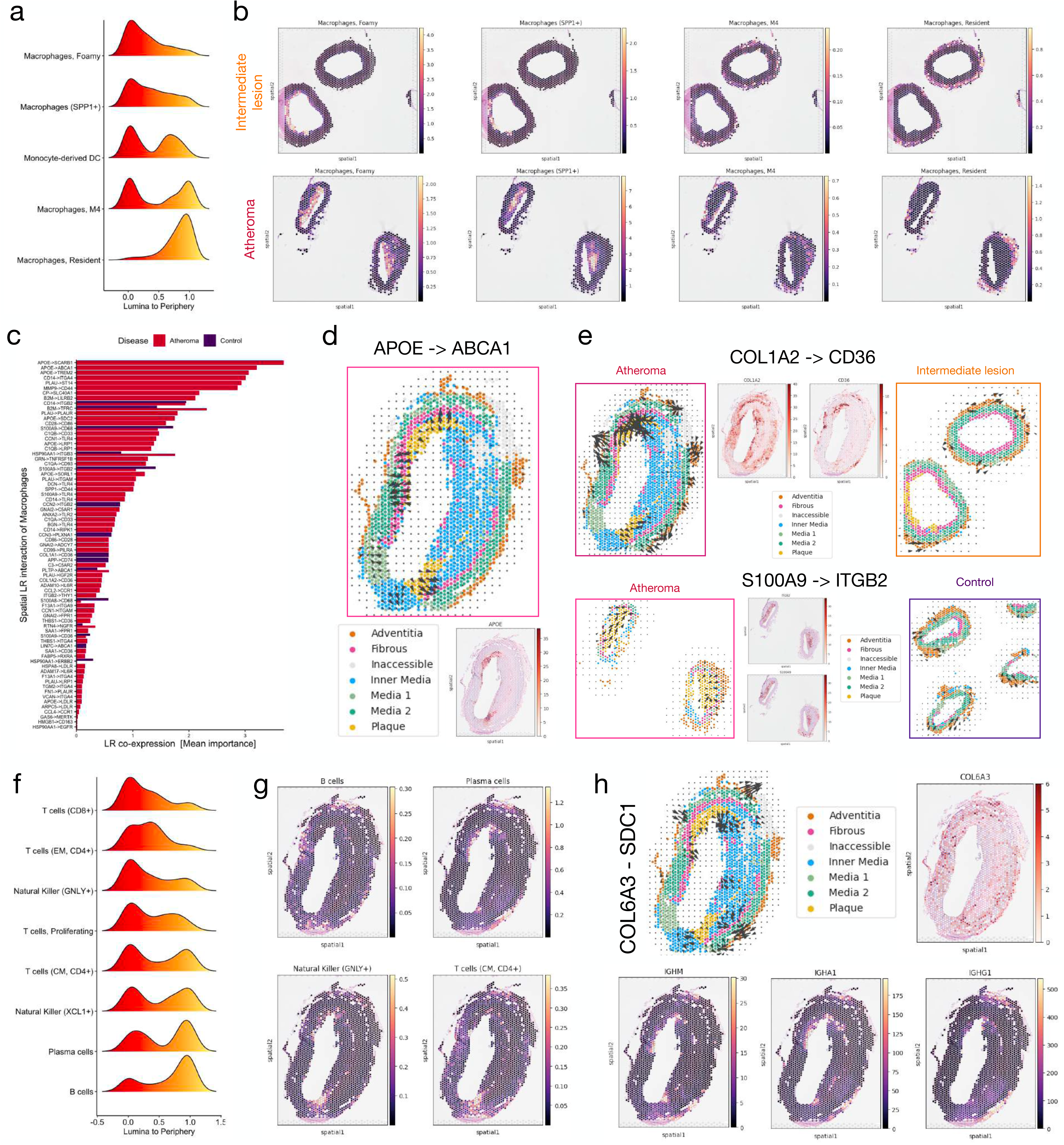
Lipid uptake, efflux and interplay of the macrophages in the human plaque. a - Ridgeline Plot demonstrating Lumina-to-Periphery scores of the relative positions of displayed macrophage subclusters within the artery across slides of intermediate lesion and atheroma. b - Depiction of cell type abundances in representative Visium slides sourced from intermediate and atherosclerotic coronary arteries, showcasing specific macrophage clusters. c - Highlighting ligand-receptor interactions highly expressed in macrophages and exhibiting high colocalization within a spot, as defined by mean importance. d - Examples of spatial Ligand-Receptor (APOE -> ABCA1) for the macrophage in the human plaque colored by the spatial microenvironment and the spatial APOE expression below. e - Spatial ligand-receptor examples for specific macrophages cell Interactions: COL1A2 -> CD36 for Spp1 positive macrophages and fibromyocytes, and also pericytes, and fibroblasts and S100A9 -> ITGB2 for M4 macrophages/neutrophils and leukocytes. The expression of the ligand and receptors in the spatial slides is shown as well. f - Ridgeline plot of the lumina to periphery score of all slides of Intermediate lesion and Atheroma displaying the relative position of the displayed lymphocyte subcluster in the artery. g - Examples of cell type abundances in representative Visium slides for the indicated lymphocytes sourced from an atherosclerotic slide. h - Spatial ligand-receptor interaction (COL6A3 -> SDC1) for plasma Cells and fibroblasts/fibromyocytes within the adventitia and toward the plaque, accompanied by spatial expression details of COL6A3 and immunoglobulin isoforms IgA/M/G1.

Subsequent spatial analysis of macrophage cell-cell communication unveiled a notable co-localization of lipoprotein APOE with ABCA1 and others (**Fig. 5c-d, Extended Data 5c**), facilitating cholesterol efflux from macrophages by reverse cholesterol transport^25^. Besides, interactions involving myeloid cells (e.g. CD33 -> C1QB) and interactions with other cell types (e.g. CCL2 -> CCR1) were depicted. Interestingly, lipid uptake driven by scavenger receptor CD36 and the interaction with COL1A1/2 were dominated by SPP1 macrophage in the plaque, but APOE pericytes in the adventitia specifically expressed CD36 resulting in COL1A2 and CD36 interaction in the plaque and adventitia (**Fig. 5e, Extended Data 5d).**

Notably, not all cell-cell interactions observed were uniquely associated with atherosclerosis, some were present in control samples. S100A9, which is specifically expressed in M4 macrophages (and Neutrophils) exhibited interaction with ITGB2 (**Fig. 5e, Extended Data 5e**), which is expressed by leukocytes associated with cytotoxicity and phagocytosis^26^. While S100A9 expression was noted in both plaque and adventitia, in atheroma slides, the interaction was predominantly localized within the plaque, contrasting with the communication observed in the control region, which was, albeit in lower presence, more inclined towards the adventitia.

In agreement with this observation, cytotoxic lymphocytes were enriched towards the lumen and the plaque peripheral border, while other lymphocytes showed a profound bimodal distribution (**Fig. 5f-g**). B cells and plasma cells showed some tendency for an adventitial localization and local interaction (SDC1 -> COL6A3) with fibroblasts and fibromyocytes (**Fig. 5h**). Interestingly, the expression of different immunoglobulin isotypes (IgA/M/G) were located proximal to the co-localizations in the plaque and adventitia, which could indicate an *in-situ* activation and plasmablast differentiation^27,28^.

Overall, the cholesterol efflux, phagocytic and cytotoxic response occur mainly in the plaque but a bimodal distribution of certain leukocytes like M4 macrophages and adaptive immune response by activated B cells and Plasma cells on the periphery might indicate an acute adventitis.

### VSMC transdifferentiation and ECM remodeling by fibromyocytes

Because of differences observed in the spatial distribution for the different pericyte subclusters and fibromyocytes (**Fig. 4e**), we next aimed to investigate it in more detail. Intriguingly, we found that pericytes (APOE+) and fibroblasts were located mainly in the adventitia, while the other pericyte cluster (MYH11+) colocalized with VSMC in the media (**Fig. 6a, Extended Data Fig. 6a**). Furthermore, we found that moving towards the lumen, the VSMC signature changed towards myofibroblasts and then fibromyocytes. Notably, fibromyocytes were detected in all different stages of atherosclerosis, including all control samples with no acute immune cell infiltration. This could indicate a general presence of fibromyocytes in human coronary arteries or them being involved in the early progression of atherosclerosis.

**Figure 6:**
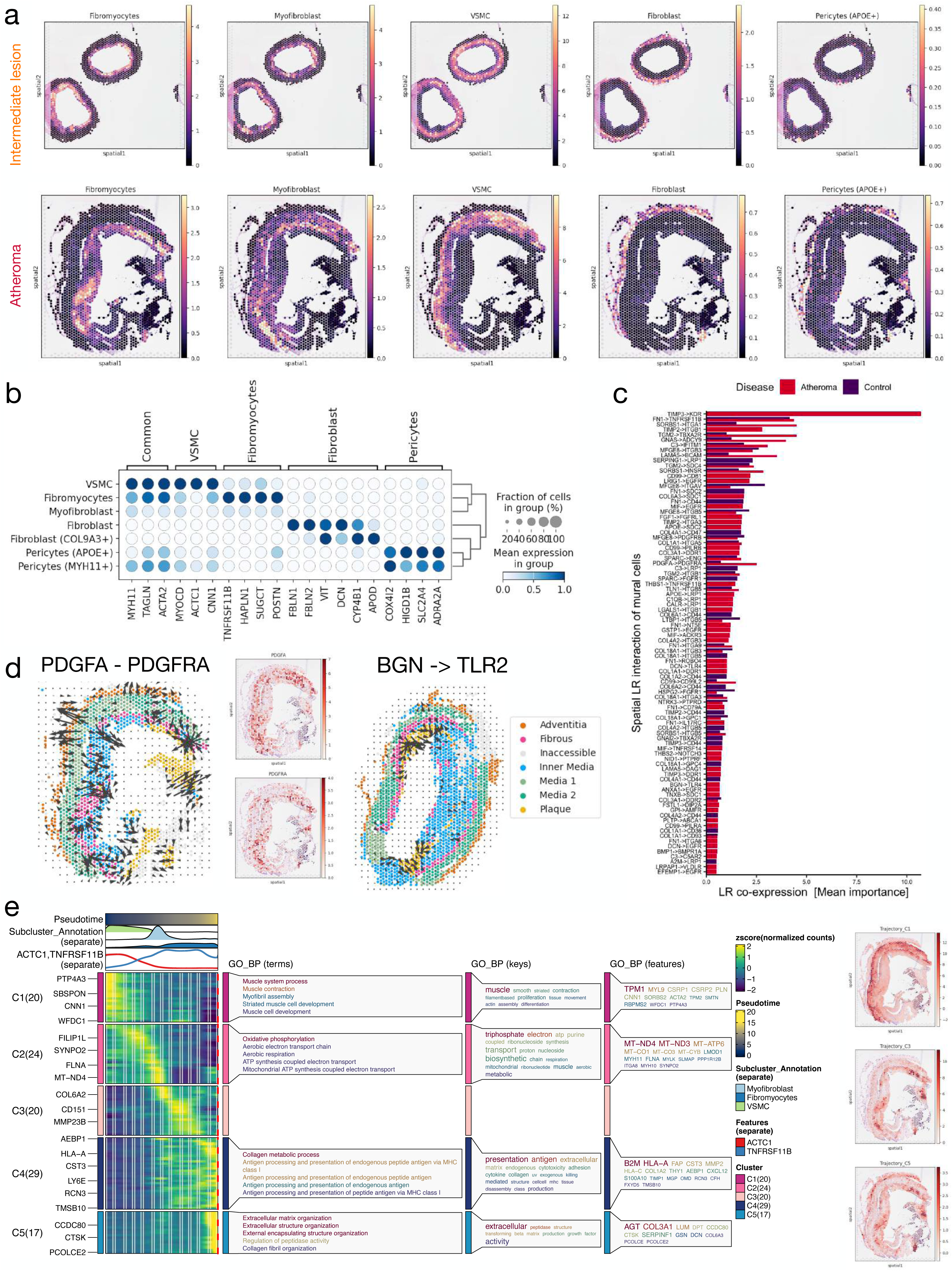
VSMC transdifferentiation and ECM remodeling by fibromyocytes. a - Visualization of the cell type abundances observed in representative Visium slides obtained from samples of both control and atherosclerotic coronary arteries. b - Identification of cell-type specific markers characterizing different cell types of VSMCs, pericytes and fibroblasts present in the human arterial wall. c - Highlighting ligand-receptor interactions expressed in fibroblasts and VSMCs, showcasing high colocalization within the spatial slides sourced from control and atheroma, as determined by mean importance. d - Examples depicting spatial ligand-receptor interactions: PDGFA -> PDGFRA interaction of VSMCs with fibromyocytes and fibroblasts and BGN -> TLR2 interaction between fibromyocytes and macrophages within the plaque core. e - Pseudotime Analysis of VSMCs: Utilization of pseudotime analysis via Slingshot to order VSMCs based on pseudotime, revealing the transition from VSMCs toward fibromyocytes via myofibroblasts. Additionally, clustering of different genes and their respective biological processes contributing to the pseudotime progression is depicted. The left side showcases aggregated expression of top genes in indicated clusters within an atheroma slide.

However, these findings together with previous studies, raised the question about appropriate markers for each cell type^29^. Historically, VSMC were defined by different markers like ACTA2, MYH11 or TAGLN and revealed the trans-differentiation diversity of VSMC, but it has been noted that MYH11 is expressed in a subset of pericytes^30^. Similarly, the analysis of the scRNA-seq atlas demonstrated expression of MYH11 for MYH11+ pericytes and all VSMC subclusters and for ACTA2 and TAGLN broadly in VSMC and pericyte subclusters. Hence, we re-analyzed the subcluster of VSMCs, pericytes and fibroblasts and suggested state specific markers for the different cell types, which alone or in combination might help to mark specific cell states (**Fig. 6b**). Notably, some of the pericyte markers were confirmed in a recent multi-organ study of pericyte markers^31^.

Next, to understand their spatial relationships, we analyzed spatial cell-cell-communication of VSMC and fibroblasts subtypes. TIMP3 and KDR/VEGFR2 colocalized highly in the adventitia of atheroma slides (**Fig. 6c, Extended Data Fig. 6b**) and is associated with angiogenesis^32^. Besides, PDGFA interactions from VSMC were observed in different directions with PDGFRA expressed in fibroblasts and fibromyocytes. Other interactions involved cell-cell communication between fibromyocytes themselves (e.g. FN1 -> TNFRSF11B) and macrophages located in the plaque (e.g. BGN -> TLR2) located in the plaque (**Fig. 6d, Extended Data Fig. 6c**).

Furthermore, the migration and transition from VMSC towards fibromyocytes has been reported previously^33^. Hence, we reasoned that the migration could occur with a cellular transition and analyzed the VSMC cluster for a potential trajectory (**Fig. 6e**). The trajectory from VSMCs via myofibroblasts and fibromyocytes occurred with expression changes from VSMC marker towards elevated expression of fibroblast markers in the end stadium (C5: DCN, COL6A3, PCOLCE), which was enriched atherosclerotic cells (**Extended Data Fig. 6d-e**). Interestingly, ECM remodeling (BGN, COL3A1, LTBP1) and antigen processing and presenting occurred along the trajectory. Finally, the defined genes from the scRNA-seq atlas displayed an increasing gradient from the media to the lumen, indicating the migration and transition of VSMC to luminal fibromyoctes. Notably, fibromyocytes in the control samples did not progress towards the later fibrous stage of fibromyocytes (**Extended Data Fig. 6f**).

Taken together, signaling from the adventitia and the plaque might regulate the communication and migration from the periphery to the lumen and fibromyocytes occur in control samples but change to a more fibroblast character upon atherosclerosis.

### Cell type and niche-specific microenvironments for drug targets

Finally, we identified bioactive compounds of the cardiovascular system according to the Anatomical Therapeutic Chemical categories with cell2drug^34^. We observed that some drugs for cardiac therapies and peripheral vasodilators (e.g. pentoxifylline) addressed different cell types of atherosclerosis, while antihypertensives (e.g. bosentan) and vasoprotectives (dexamethasone acetate) acted mostly in pericytes and macrophages respectively, and beta blockers or calcium channel blockers had a high association with Mast cells (**Fig. 7a, Extended Data 7a**). Drugs of the renin-angiotensin system acted broadly on atherosclerotic cells (e.g. lisinopril) or in a more cell-type specific manner (e.g. losartan on pericytes) in agreement with their different mechanisms, while lipid lowering agents such a statins (e.g. lovastatin, simvastatin) were targeted towards macrophages. Interestingly, the observed cell-type specificity or broadness of the drugs could be recapitulated on the spatial transcriptomic slides (**Fig. 7b**). And while statins showed the expected action in the core plaque region, Losartan acts on the periphery and thus highlights the relevance of the microvasculature for atherosclerosis treatment.

**Figure 7:**
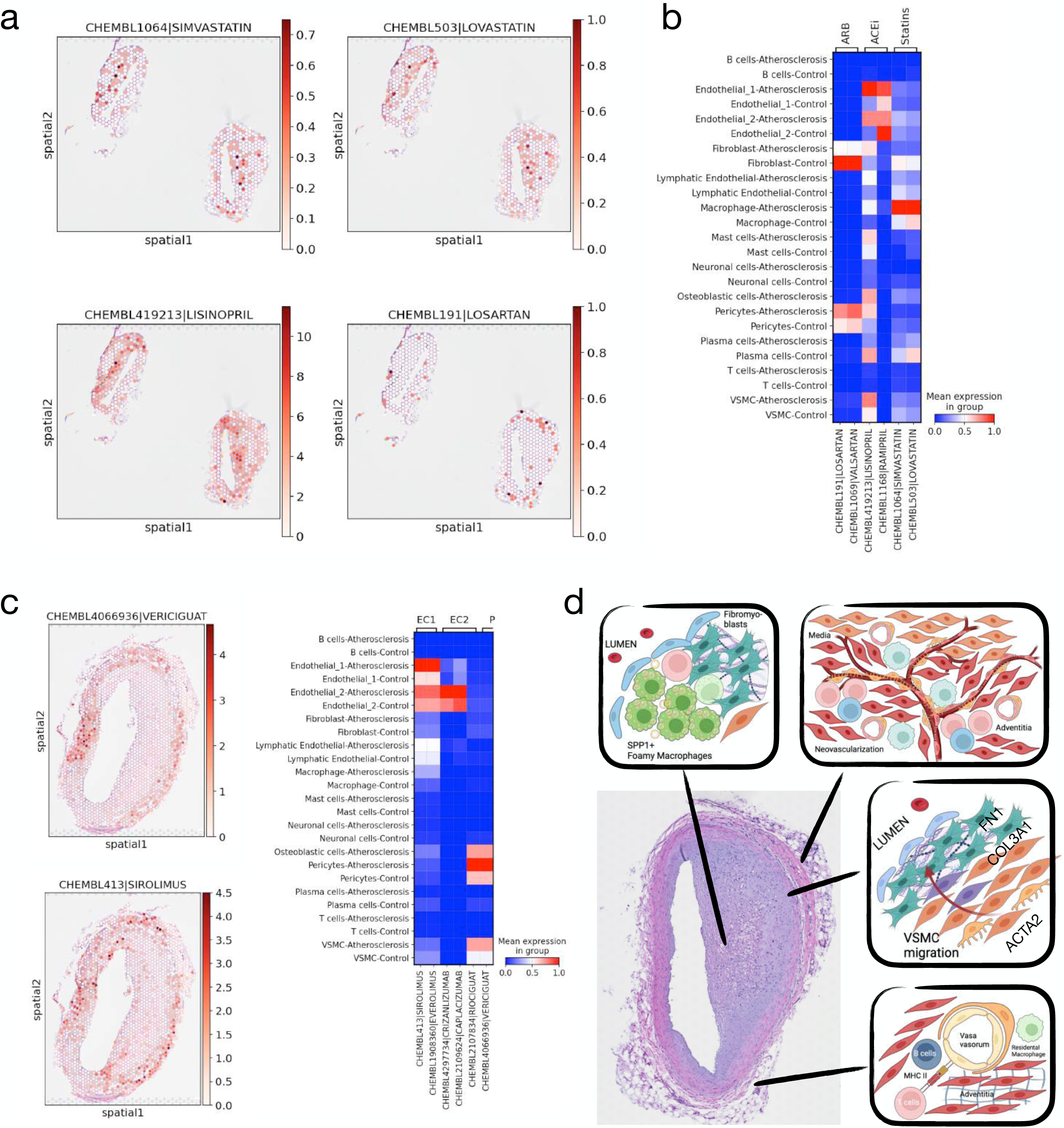
Cell type and niche-specific microenvironments for drug targets. a - Drug target expression of ChEMBL bioreactive compounds for different drugs that are used for atherosclerosis treatment. b - Visual representation of spatial target expression in an exemplary atheroma slide specifically focusing on ACE inhibitors (ACEi), Angiotensin Receptor Blockers (ARB), and statins. c - Potential drug repurposing candidates with specificity for peripheral endothelial cells and macrophages (Sirolimus), luminal endothelial cells (Crizanlizumab), or pericytes (Vericiguat). Spatial expression patterns of these drug targets are illustrated in a single slide (left) and in the scRNA-seq data of the human atherosclerosis atlas (right). d - Illustrating key findings related to various niches within the human arterial wall, including macrophage interactions and lipid efflux within the plaque, neovascularization in the adventitia, VSMC migration and differentiation toward the lumen to become fibromyocytes, and recruitment of lymphocytes from the vasa vasorum with adhesion to local endothelial cells and pericytes.

Next, we analyzed the human atherosclerosis atlas for cell-type specific drug targets. We observed differences for potential drug targets in the two endothelial cell clusters, where adventitial endothelial might be the potential target of Rapamycin-like targets (e.g. Sirolimus) which could inhibit broadly cell proliferation of endothelial and immune cells^35^, while Crizanlizumab/Caplacizumab might be specific drugs for targeting luminal endothelial and immune cell interactions (**Fig. 7c, Extended Data 7b**). Potential drug targets for pericytes were widely classified for the cardiovascular system, which included vericiguat, a stimulator of the soluble guanylyl cyclase (GUCY1A1) that has been recently suggested as a potential atherosclerosis treatment^36^.

Overall, we found specificity of existing drugs for certain cell types and the scope of action. Novel targeting approaches might benefit from taking cell type specificity and/or the spatial niche and tackling specific interaction in an atherosclerotic artery.

## Discussion

The cellular composition in atherosclerosis has previously been studied by exploration of circulating blood cells or predefined markers applied to the human artery^37^. Using single-cell RNA-seq and spatial transcriptomics, the cellular heterogeneity can be explored in an unbiased manner^38^. Furthermore, by integrating multiple studies, deeper insights from a more encompassing view in human atherosclerosis can be drawn from the extended cohorts^39^.

Our integrated scRNA-seq atlas illuminates an increased proportion of neuronal cells within coronary arteries and an intriguing prevalence of osteoblastic cells within the carotid arteries. In agreement, connecting work on the peripheral nervous system demonstrated axogenesis in the adventitia^40^, while the osteoblastic cluster has not been described in individual studies. Osteoblastic cells, consistent of carotid arteries from different patients, expressed marker for VSMC (ACTC1, MYH10), osteoblast differentiation (DLX5/6) and Sclerostin (SOST), a WNT-inhibitor and regulator of bone mineralization with athero-protective features^41^. However, despite the integration of multiple studies and consistency of carotid arteries from different patients to the osteoblastic cluster, the definitive specificity to carotid atherosclerosis remains elusive due to controls of the scRNA-seq atlas originating from coronary arteries.

Furthermore, our atlas suggests pericytes to be involved in leukocyte attraction, neovascularization and hemorrhage in atherosclerosis as evidenced by their changes in expression. The role of pericytes in immune modulation^42^, angiogenesis^43^ and differentiation^44^ has been studied in different tissues and rendered them interesting as therapeutic targets in cardiovascular diseases^45^. However, their expression state is very heterogeneous and variable within a tissue and between tissues^46^. Hence, we suggested markers of the scRNA-seq atlas specifically expressed in vascular pericytes and widely in agreement with a multi-organ study of pericytes^31^.

Apart from that, two different endothelial cells were identified from our scRNA-seq atlas, which by the addition of our spatial transcriptomics data of human coronary arteries, could be spatially assigned to lumina and peripheral endothelial cells. Furthermore, the spatial data revealed further spatial distribution and local cellular communication in an unbiased approach (**Fig. 7d**).

In agreement with previous studies, we demonstrated the accumulation of lymphocytes in the adventitia^47,48^. However, with the unbiased approach of spatial transcriptomics, we analyzed their potential interactions with adventitial endothelial cells and pericytes and found that the Major Histocompatibility Complex II, which can support the antigen-presentation and facilitate the adaptive immune response^21,49^ and the expression of ACKR1, which might be involved in angiogenesis, chemotaxis, and cellular retention signals^50^ were higher expressed in adventitial than in luminal endothelium. Notably, pericytes have also been demonstrated as antigen-presenting and cells for T-lymphocytes^43^. While the question remains elusive if adventitial cells might act as antigen-presenting cells, the presence of immunoglobulin-producing plasma cells might indicate activation of B cells^27,28^ and an acute state of inflammation of the adventitia, termed adventitis^47^.

While the accumulation of T-cells and especially macrophages is highest in the plaque, important factors of the cholesterol and lipid intake and efflux (e.g. CD36, APOE, ABCA1) showed a high transcription in this location. Interestingly, CD36 which has been reported to be important for lipid uptake into VSMC and other cells^51^ have been found to be expressed in pericytes but not VSMC. While the context might need to be evaluated^29,30^, CD36 might regulate the angiogenesis by inhibition of VEGFR2/KDR^52^.

Investigation of integrated datasets has unraveled the dynamics of modulated SMC phenotypes in atherosclerosis^33^. We elaborated on it and demonstrated that VSMC can act as a signal mediator to luminal fibromyocytes as well as adventitial fibroblasts and we showed a spatial trajectory indicating migration and transition of VSMC into fibromyocytes and in the case of atherosclerosis a progression toward a fibroblast-like expression state of the fibromyocytes.

The different aspects of this work could indicate different drug target approaches, including cell type specific targets, regulation of the immune cell infiltration, lipid efflux stimulation or ECM remodeling. Here, several drugs for the cardiovascular, especially antihypertensives addressed targets in pericytes, consistent with their regulation of blood pressure^53,54^.

We acknowledge that inherent limitation of lacking genetic data precludes the definitive establishment of causality within the scope of our study. Further limitations of this study stemming from the sparsity of single-cell transcriptomics data, and inherent technical batch effects from unknown sources. Furthermore, some of the cell-states identified in a computational analysis need to be carefully experimentally validated. Nonetheless, our robust analyses, inclusion of multiple studies and integration with novel spatial data present a valuable resource for understanding atherosclerosis and further investigations how drugs interact with specific cell populations in distinct spatial locations within the artery and hence opens new horizons for precisely tailored treatments.

In summary, our results underscored the impact of spatial resolution in the human artery and emphasizes the need for further research of the role of the microvasculature in plaque development. Importantly, the role of acute adventitis in atherosclerosis warrants detailed investigation, potentially serving as a supportive side-effect, presenting contradictory phenomena, or even playing a partial initial causative role.

## EXTENDED DATA FIGURES

**Extended Data Figure 1:**
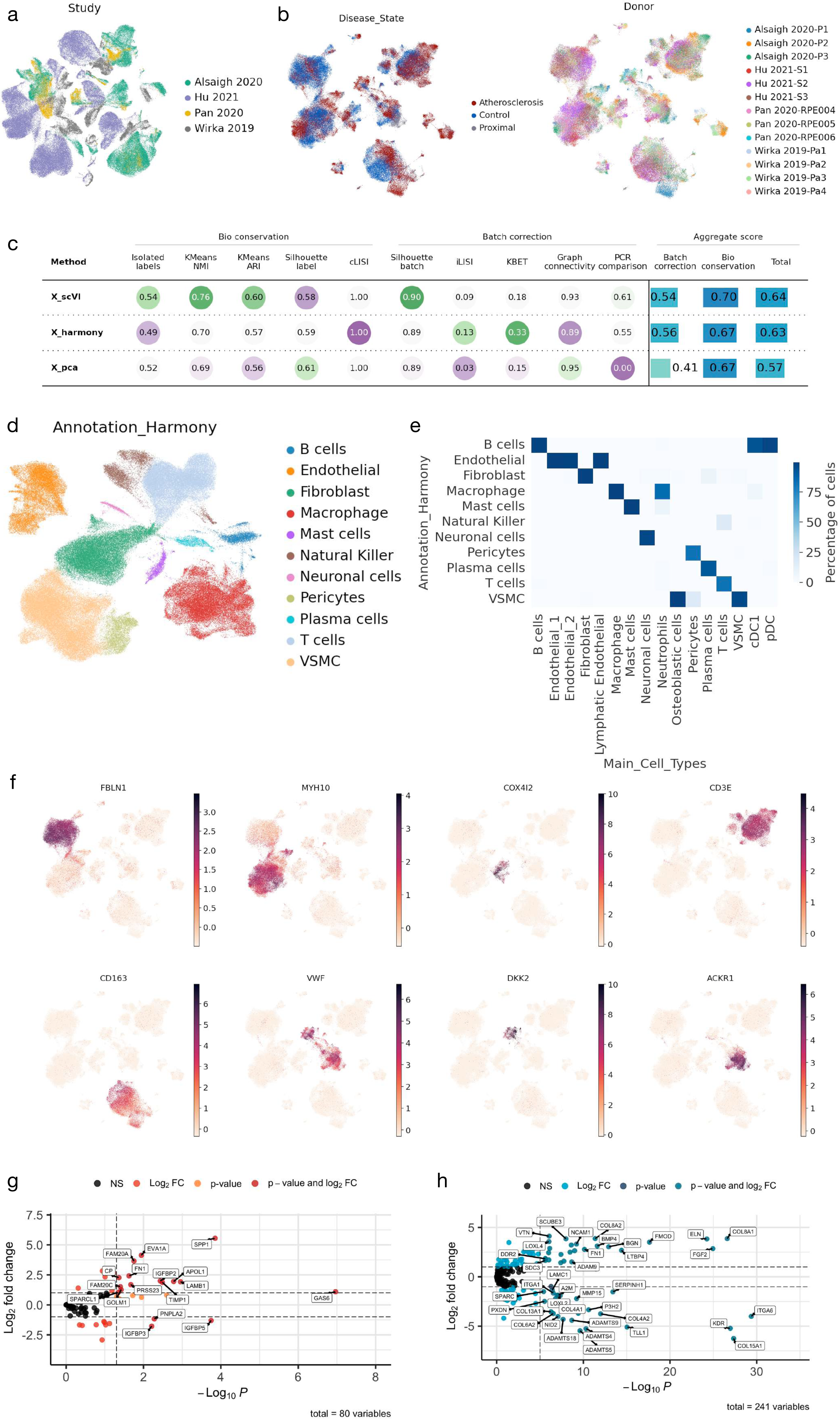
a - UMAP of the unintegrated sc RNA-seq data sets colored by the studies. b - UMAP of the scVI-integrated scRNA-RNA seq data colored by the atherosclerosis state and the donors. c - Benchmarking results of the different integration approaches using scIB. d - UMAP of the Harmony integrated data set colored by the cell types that have been annotated by marker gene expression. e - Comparison of the cell type annotation based on the integration and clustering with scVI (x-axis) and Harmony (y-axis). The color indicates the percentage of overlapping single-cells for each cluster annotation. f - Examples of marker gene expression for the scVI-integrated human single-cell atherosclerosis atlas. fibroblasts (FBLN1), VSMCs (MYH10), pericytes (COX4I2), T-cells (CD3E), macrophages (CD163), endothelial (VWF) and specific marker for Endothelial_1 (ACKR1) and Endothelial_2 (DKK2) are shown. g - Volcano plot of the differential expressed genes of the Reactome pathway REGULATION_OF_INSULIN_LIKE_GROWTH_FACTOR_IGF_TRANSPORT_AND_U PTAKE_BY_INSULIN_LIKE_GROWTH_FACTOR_BINDING_PROTEINS_IGFBPS between control and atherosclerosis of the pericytes cell type cluster. h - Volcano plot of the differential expressed genes of the Reactome pathway EXTRACELLULAR_MATRIX_ORGANIZATION between the cell type cluster Endothelial_1 and Endothelial_2.

**Extended Data Figure 2:**
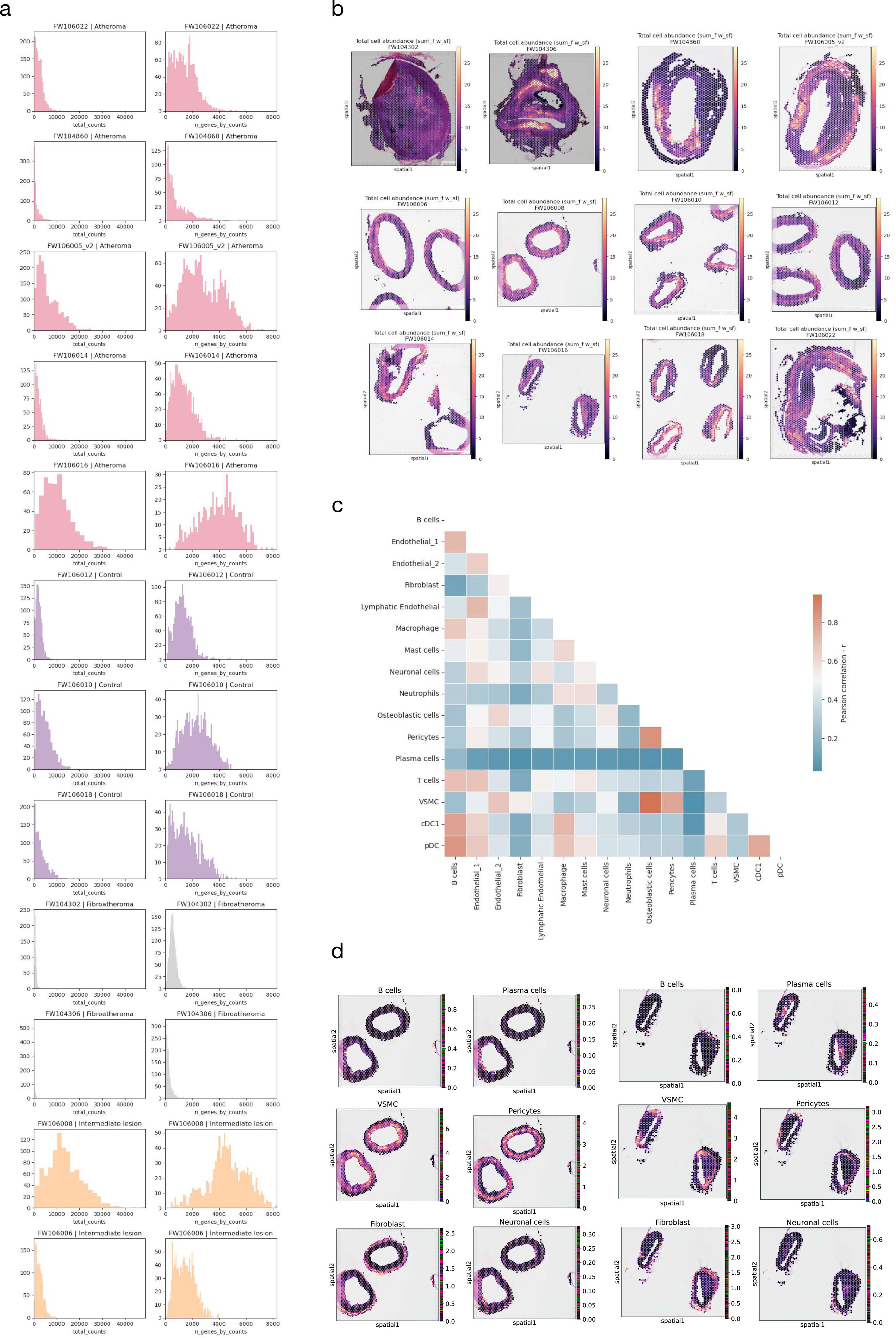
a - Quality Control of the Visium slides showing the total counts and the n_genes_by_counts for each specimen colored by the atherosclerosis state. b - Overview of the Visium slides and their total cell abundance. c - Pearson correlation of the reference signature of the scRNA-seq human atherosclerosis atlas that overlapped with the genes detected in the Spatial slides. d - Examples of cell type abundances in Visium slides from a specimen with intermediate lesion or atheroma of the coronary artery.

**Extended Data Figure 3:**
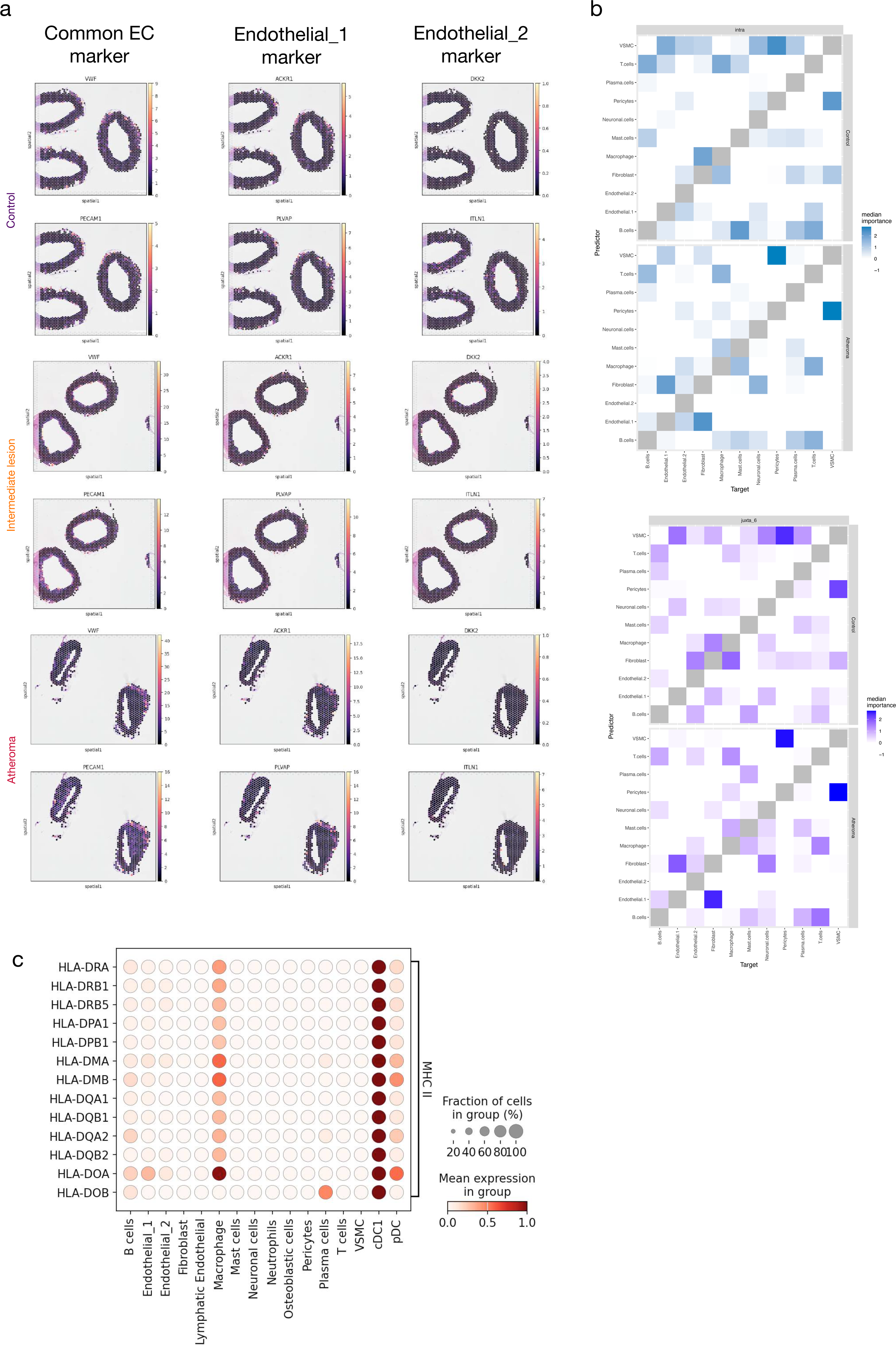
a - Examples of different endothelial marker gene expression for the representative control, intermediate lesions and atheroma slides. b - Misty results for the co-localization of the intra and proximal view of the control and atheroma slides. c - Expression profile of the Major Histocompatibility Complex (MHC-II) in the human atherosclerosis single-cell atlas.

**Extended Data Figure 4:**
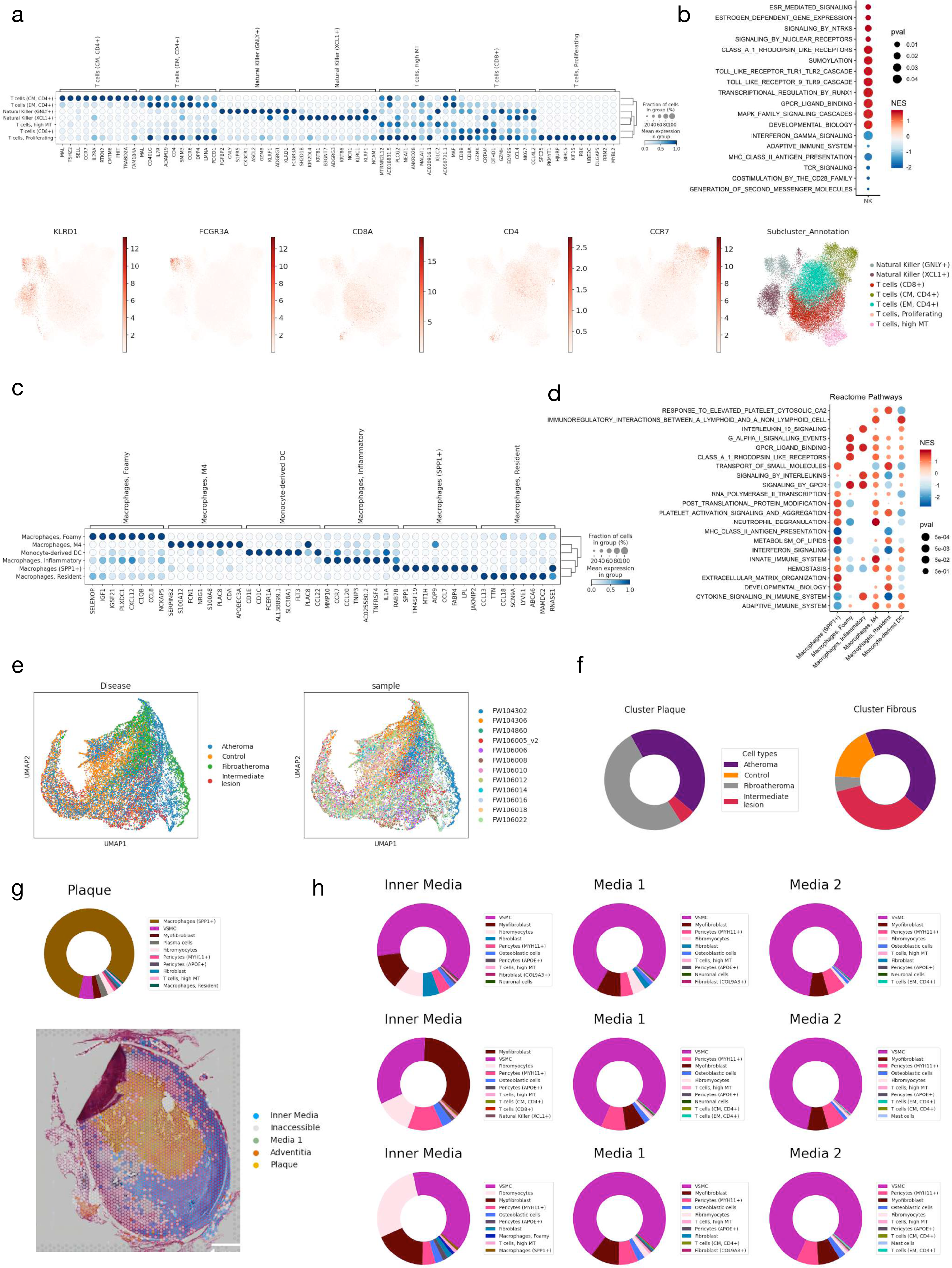
a - Top marker for each sub cluster cell type of the T cell cluster of the human atherosclerosis scRNA-seq atlas. Below the UMAP of the T cell subclustering and selected markers for the cell type identification are shown. b - Differentially expressed genes in the two Natural Killer cell clusters and their corresponding reactome pathway enrichment for differential expression between the clusters. The color represents the normalized enrichment score (NES) and the p-value from the gene set enrichment analysis is represented by the size. c - Top marker for each sub cluster cell type of the macrophage cluster of the human atherosclerosis scRNA-seq atlas. d - Differentially expressed genes in the different macrophage subclusters and their corresponding reactome pathway enrichment for differential expression between the clusters. The color represents the normalized enrichment score (NES) and the p-value from the gene set enrichment analysis is represented by the size. e - UMAP of the Visium spot clustering colored by the atherosclerosis state and the specimen. f - Proportions of the disease progression background highlighted for the plaque and fibrous cluster. g - Proportion of the plaque cluster for the most advanced atheroma slides and an example with the spatial location of the clusters. h - Median cell type abundance within the spots of each spatial microenvironment indicated for the sample from control, intermediate and atherosclerotic coronary arteries.

**Extended Data Figure 5:**
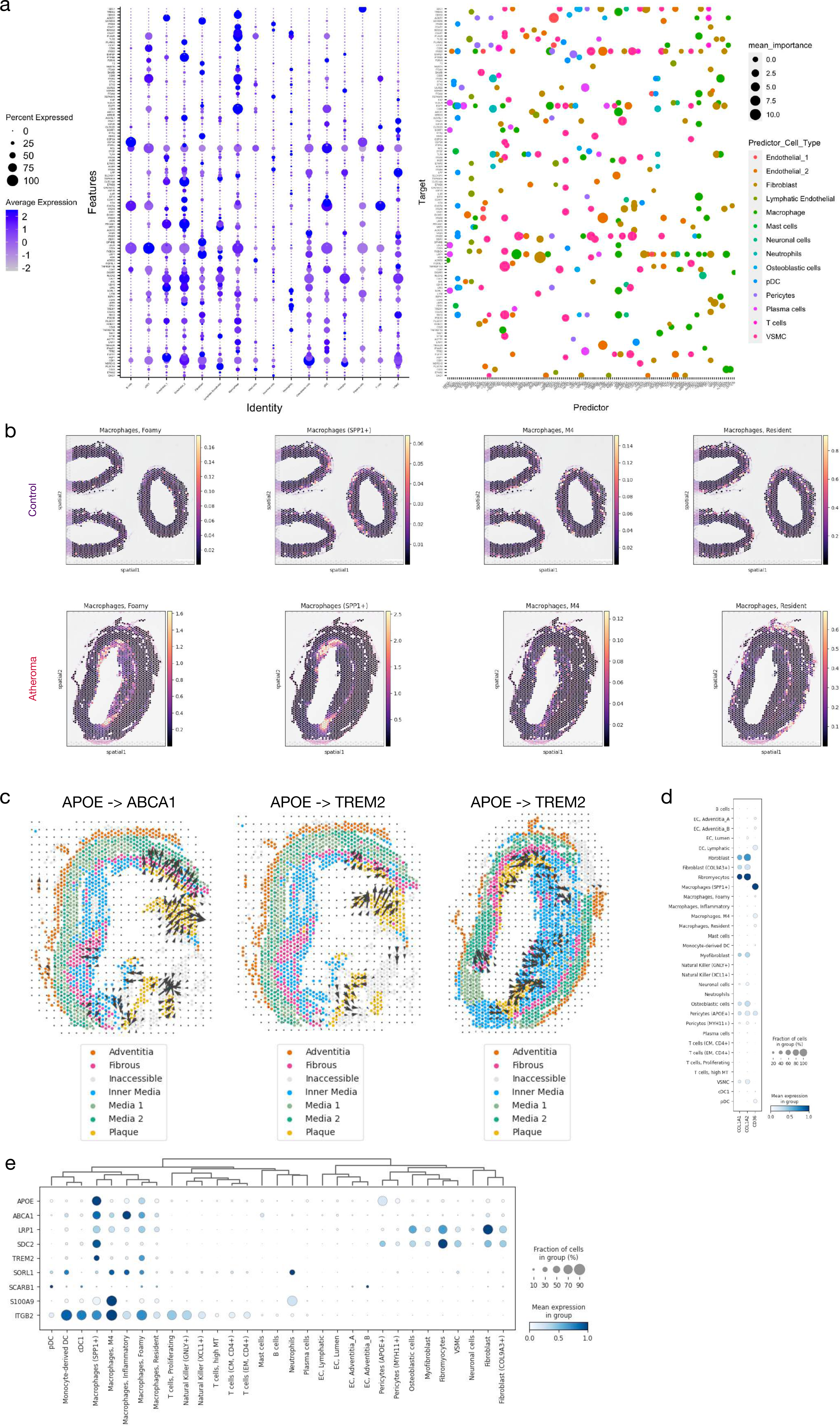
a - Overview of the spatial relevant ligand-receptor interaction in atherosclerosis. On the right the receptor expression in the atherosclerosis atlas is ordered by cell type specific expression and on the left the co-localization as mean importance of all atheroma slides with the cell type of the ligand (predictor) colored and ordered by cell type specificity. b - Examples of macrophage subcluster cell type abundances in Visium slides from control and atherosclerosis coronary artery. c - Examples of spatial Ligand-Receptor (APOE -> ABCA1 and APOE -> TREM2) for the SPP1+ macrophages / Foamy macrophages of the plaque core colored by the spatial microenvironment. d - Expression profile of the Receptor-Ligand COL1A1/2 and CD36 in the human atherosclerosis atlas. e - Expression profile of the Receptor-Ligand involved macrophages and their expression in the human atherosclerosis single-cell atlas.

**Extended Data Figure 6:**
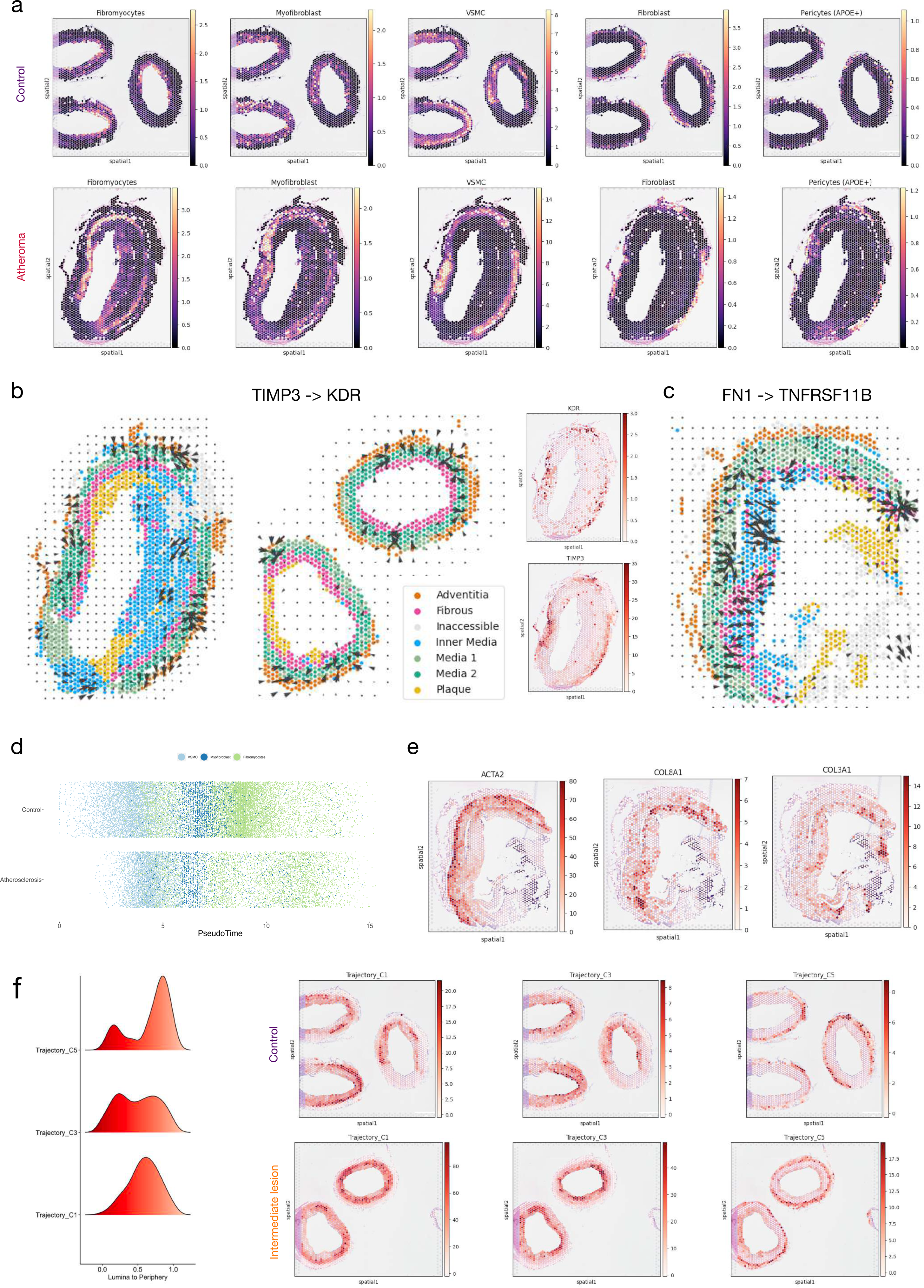
a - Examples of different subcluster cell type abundances in Visium slides from control and atherosclerosis coronary artery. b - Examples of spatial Ligand-Receptor (TIMP3 -> KDR) for the fibroblasts and adventitial EC colored by the spatial microenvironment. c - Examples of spatial Ligand-Receptor (FN1 -> TNFRSF11B) for the intra-fibromyocytes interaction. d - Individual VSMCs of the scRNA-seq atlas ordered by the Pseudotime of the trajectory analysis and colored by their subcluster level. e - Example expression of Trajectory Cluster gene expression in an atheroma slide with ACTA2 (C1), COL8A1 (C2) and COL3A1 (C3). f - Density plot of the cluster accumulation in the atheroma slide shown in Figure 6 and two examples of the Trajectory cluster expression in control and intermediate lesion.

**Extended Data Figure 7:**
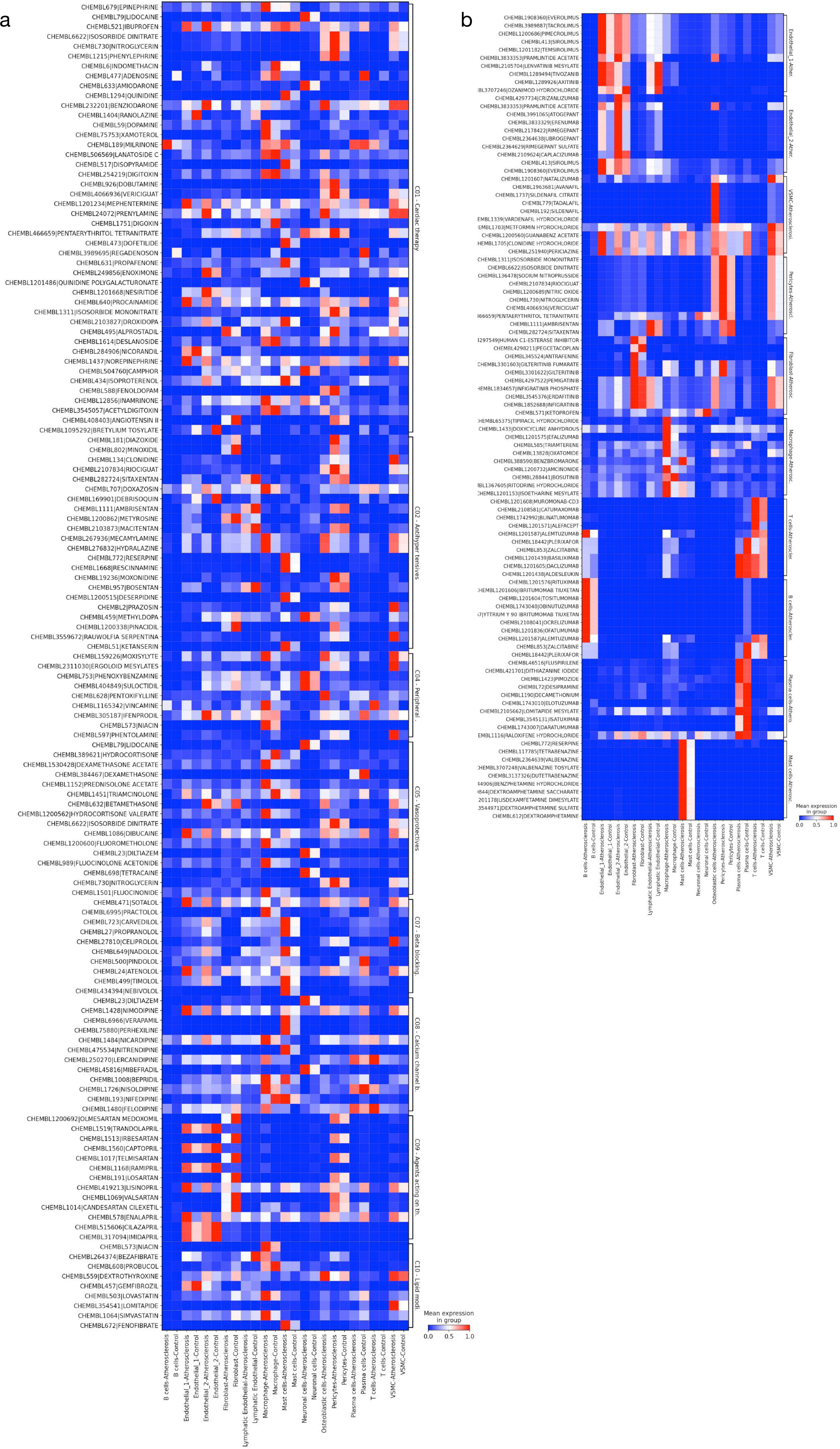
a - Drug target expression of ChEMBL bioreactive compounds for the Cardiovascular system and the different ATC categories. b - Drug target expression for putative cell-type specific ChEMBL bioreactive compounds for the major cell types of human arteries.

## Data availability

The processed single-cell RNA-seq atlas and the 10X Visium slides of the human coronary arteries will be available at GEO with the publication of the manuscript.

## Code availability

The code is available at Github: https://github.com/hayatlab/multiomics_athero_atlas

## Acknowledgments

We wish to thank laboratory technicians Heidi Solvang Nielsen and Jeanette Juul (both from Novo Nordisk A/S) for skillful technical assistance with the wetlab work in the Visium workflow.

## Contributions

TB, VD, SB, RK, SH developed the concept, HH, LMV, CP generated Visium data, SM performed CrosstalkR analysis, TB, AB, MH annotated cell cluster, TB analyzed, interpreted data and wrote the manuscript with inputs from RK and SH. Supervision and funding Acquisition was conducted by VD, SB, RK, SH. All authors reviewed the manuscript, helped with interpretation of the results and revised the manuscript.

## Methods

### Data collection and preprocessing

For the single-cell atlas, public available single-cell RNA-seq data from carotid and coronary arteries were searched. The data sets from all studies were produced with the 10X Genomics technology by the authors and were obtained either directly in the counts of GRCh38 Ensemble v93 genes or re-analyzed with cellranger v3.1.0 and GRCh38 Ensemble v93 genes to obtain the same gene annotations for all data sets. For each data set, cells with a minimum of 200 features, a maximum of 20 % mitochondrial reads and a doublet rate < 3 determined by scDblFinder^55^ were considered.

### Single-cell data integration and cell type annotations

For the single-cell data integration, 8,000 highly variable genes were selected for each sample (flavor="seurat_v3", batch_key="Donor") and Harmony^12^ with default settings and scVI^13^ with 2 layers and 30 latent dimensions were used. For the benchmarking of the integration methods, the scIB benchmarking framework was used^56^.

For the main cell type annotation, different clustering resolutions of the integrated embedding were evaluated. Leiden Clustering with a resolution of 0.2 was chosen for the main cell type cluster annotation. The main clusters were independently sub-clustered at different resolutions accordingly and the optimal sub-clustering was chosen based on marker genes and manual inspection.

### Differential expression and pathway analysis

Differentially expression was determined with the scVI differential expression module in change mode and enabled batch correction. The genes were further filtered by bayes factor >2 and non_zeros_proportion > 0.05. Independently, differentially expressed genes were determined from raw counts for pseudo-bulk analysis per sample per cluster and analyzed with DESeq2 (Version 1.38.3) and p-values < 0.05.

Fgsea 1.24.0 was used to perform pathway analyses^57^ with differentially expressed genes (FDR < 0.08) obtained from the scVI DE module between disease conditions or cell types. Reactome gene sets (with the number of genes > 15 and < 500) were used. Reactome pathways were obtained from msigdb database (Version 7.5.1: https://bioconductor.org/packages/msigdb). Only the pathways with a p-value < 0.05 for a given cell-type are reported.

### Single-cell RNA-seq Cell-Cell-Communication

CrossTalkeR package (v. 1.3.6) with a liana wrapper for the LR pairs were used to predict ligand receptor (LR) interactions between different cell types in the scRNA-seq atlas. The *liana_wrap* function of liana (v. 0.1.12) calculates the LR prediction on the basis of different methods as natmi, cellphonedb, and liana. From this we used the cellphoneDB predictions for the input of CrossTalkR to generate a comparative cell-cell interaction (CCI) network for different conditions. Then for the CCI analysis of the immune cells and endothelial cells / pericytes specific interactions that occurred in less than 40 interactions were considered.

### Spatial transcriptomics

Sections of 5 µm thickness were cut from formalin-fixed and paraffin-embedded (FFPE) samples of human coronary arteries and mounted on Visium slides and processed for spatial transcriptomics according to the 10X Genomics Visium FFPE Version 1 protocol. Briefly, samples were deparaffinized, stained with hematoxylin and eosin (H&E) and imaged using VS200 Slide Scanner (Olympus Life Science) prior to decrosslinking, destaining and overnight probe hybridization with the 10X Visium Human version 1 probe set. The following day, hybridized probes were released from the tissue and ligated to spatially barcoded oligonucleotides on the Visium Gene expression slide. Barcoded ligation products were then amplified and used for construction of libraries, which subsequently were sequenced on a NovaSeq 6000 sequencing platform (Illumina), using a NovaSeq 6000 S2 Reagent Kit v1.5 (Illumina) according to the manufacturer’s instructions. With SpaceRanger version 1.3.0 (10X Genomics) reads were aligned to their corresponding probe-sequences (Visium human transcriptome probe set v1, based on GRCh38 2020-A) and mapped back to the Visium spot where a given probe were originally captured, and finally aligned to the original H&E-stained image of the tissue section. The filtered count matrix .h5 file was used for further downstream processing and analysis of data.

### Spot deconvolution and atherosclerotic microenvironment

Using our scRNA-seq atlas with different cell type annotation levels, the cell states were mapped to the Visium data by cell2location^18^. First we excluded genes and cells with a cell_count_cutoff=5, cell_percentage_cutoff2=0.03 to estimate the reference signatures of cell states from the scRNA-seq with a negative binomial regression model provided in the cell2ocation package. With the interfered reference signature from the spatial slides, cell2location was used to estimate the abundance of each cell type in each Visium spot with an estimate of 5 to 10 cells per spot and detection_alpha=200 as hyperparameter.

For the spot clustering we used the cell type deconvolution to calculate the neighbors and Leiden clustering (resolution = 0.2), manually annotated the resulting clusters according to their location in the artery and calculated the absolute values and cell type proportions for each cluster.

### Spatial distance metrics and co-localization analyses

For the lumina_2_periphery score, we computationally determined the center of the lumen with an optional offset based on its shape. From there the closest spots of the luminal layer and the farest spots in the periphery for 90 sectors were determined. All spots in between were assigned values of 0 (lumina) to 1 (periphery) based on their spatial location.

MistyR (version 1.8.1) ^19^ was used to estimate the importance of the abundance of each main cell type in explaining the abundance of the other cell types by modeling cell-type cell2location estimations of the control and atheroma slides. Here, an intrinsic view was applied to measure the relationships between the deconvolution estimations within a spot, and a juxta view was used to sum the observed deconvolution estimations of immediate neighbors (largest distance threshold = 6 spots). The aggregated estimated standardized median importances of each view of the slides were interpreted as cell-type dependencies in different spatial contexts, such as colocalization or mutual exclusion.

### Spatial DE and Cell-Cell-Communication

For the spatial Cell-Cell communication analysis, the slides were filtered for receptors and ligands from the LIANA database with an adequate expression of at least 250 counts per slide. Then MistyR (version 1.8.1)^19^ was used to determine the importance of the intrinsic view within a spot and the juxtaposition of the surrounding 6 spots. The mean importances were calculated per disease state and view. Predictor and Target matrix was filtered by existing interactions in the LIANA data base and cell type were assigned to ligand and receptors based on the highest average expression in the respective cluster in the scRNA-seq atlas. The cell type specificity determined the specificity of the assignment of this cell type by calculating the ratio to the next highest average expression. Based on the cell type assignment, cell-cell interactions were further processed. Spatial differential expressions were estimated on bivariate Moran’s R and for the cell-cell communication interplay, the LIANA ligand-receptor database was subjected to COMMunication analysis by Optimal Transport (COMMOT)^22^ with a spatial distance of 400 µm.

### Trajectory analyses and cell2drug

Trajectory analysis of selected clusters was performed with the scVI embedding and the DynamicHeatmap function of the single-cell pipeline (https://github.com/zhanghao-njmu/SCP) and 5 clusters. The top genes obtained for each cluster were aggregated with the scanpy score_genes function on the visium slides.

Drug2cell^34^ was applied to identify the Chembl listed bioactive molecules for the Anatomical Therapeutic Chemical (ATC) classification of the cardiovascular system or by differential expression analysis of the drug2cell assay considering only cells from atherosclerosis for the differential expression analysis.

